# Intermittent Cranial Accelerometric Pulses - a Potential Marker of Neurodegenerative Disease

**DOI:** 10.64898/2026.08.28.747878

**Authors:** Christine M. Walsh, Paul A. Lovoi, Leslie Yack, Joseph Chen, Natalie Pandher, Eliot D. Lee, Thomas Hemphill, Esther Li, Dominica Randazzo, Steven H. Woodward, Thomas C. Neylan, Wade S. Smith

## Abstract

We identified Intermittent Cranial Accelerometric Pulses (ICAPs) also known as “Sleep Bursts” (SBs), as a novel phenomenon of brief (1-2 sec), often periodic bursts in cranial forces occurring during human sleep. Our goal was to characterize SBs in normal subjects, then compare SB in a cohort of subjects with neurodegenerative disease (NDD). We recorded 32 cognitively healthy subjects (23-87 years) and 13 subjects with NDD (51 – 84 years). SBs occurred in all 45 subjects. SBs occurred at 0.57 SB/min (once per 105 seconds) in controls and 0.40 SB/min (once per 150 seconds) in NDD (p = 0.0043). SB occurred with equal rates across all sleep stages in both groups. When occurring periodically, SBs had modal intervals (3.75 bursts/min (0.0625 Hz) - 2.67 bursts/min (0.044 Hz)). EEG power increased in the delta range 1-2 seconds before and following the SB. EEG delta power during a SB was significantly lower in all NDD subjects across sleep stages compared to controls. The relatively low frequency of SB events and synchronization with EEG power has no parallel in human sleep; we hypothesize that SBs may represent a brain-generated pulsatile component of brain glymphatic drainage.

## Introduction

The reasons mammals sleep are enigmatic. Sleep appears to restore brain functions to normal and is involved in memory consolidation and motor learning(*1-4*). Sleep deprivation is harmful to humans, and when chronic, is associated with increased risk for diabetes, cardiovascular disease, cognitive impairment, accidents and injuries, dementia, and all-cause mortality(*5, 6*). Though sleep is important for human brain and body health, it remains less clear how sleep physically manipulates and modulates human physiology. This is mostly due to limited tools for measuring human physiological measures without perturbing naturalistic sleep. One example of a system that seems intrinsically associated with sleep is the “glymphatic system”, which flushes proteins and metabolites into the interstitial spaces of the brain during sleep, in a pulsatile manner, and is a candidate mechanism for supporting sleep’s restorative role(*7, 8*). The sense of feeling well-rested may be the consequence of adequate nocturnal cleansing of brain metabolic waste. Further, impaired glymphatic flow has been hypothesized as enhancing the concentration of brain proteins found in Alzheimer’s disease and other pathological protein-aggregate diseases(*7, 9, 10*).

### Statement of Specific Scope

We report the novel observation that humans experience short-duration, often periodic, increases in cranial forces emerging from inside the head during normal sleep (Figures 1 and S2).

**Figure 1.**
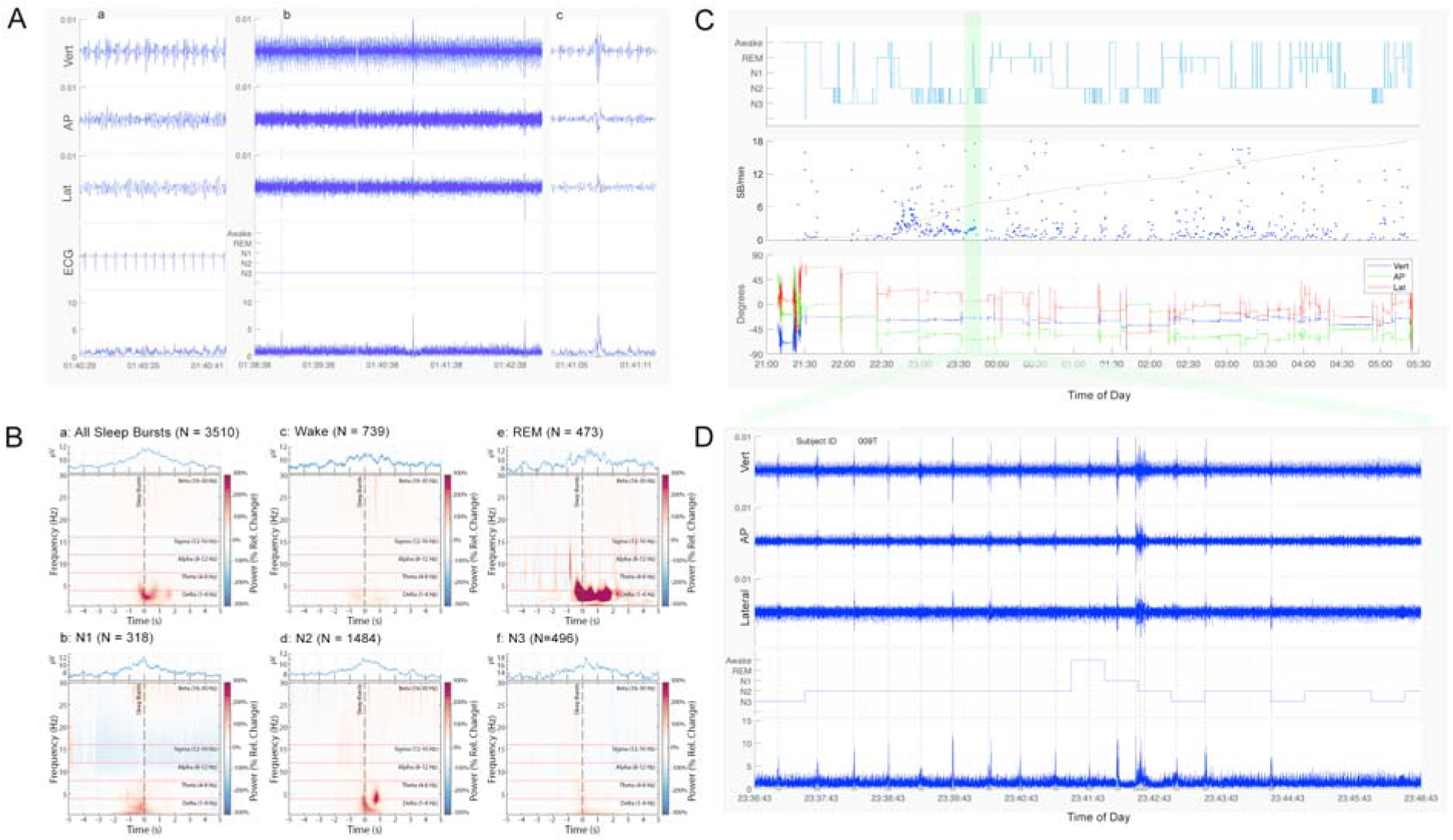
(**A**) Example of the headpulse and Sleep Bursts (SBs). (**Aa**) Headpulse in the vertical (Vert, top), anterior posterior (AP, second from top), and lateral planes (Lat, third from top). The ECG is shown second from the bottom. The bottom trace is derived from a combination of all three axes and rectified to enhance transient increases in forces from the background (sleep burst trace defined in methods). (**Ab**) Example of three SBs occurring 126 sec and 102 sec apart. The vertical dotted line indicates the timing of the center of the SB. (**Ac**) Detail of the third SB from panel b. The transient increase in power shown in the bottom trace lasts about 3 cardiac cycles in duration. (**B**) Frontal Electroencephalography Time Frequency Representation of Sleep Burst Events. The time-frequency representation (TFR) of the frontal EEG (Fpz) is shown from −5 s to 5 s relative to sleep burst events with warm colors (red) denoting spectral power increases and cool colors (blue) denoting spectral power decreases relative to the baseline of −5 to −4 s relative to sleep bursts. The rectified, averaged, EEG trace is included inset above the TFR averaged across sleep bursts in (A) all stages (N=3510), (B) wake stages (N=739), (C) rapid eye movement stages (REM, N = 473), (D) N1 stages (N = 318), (E) N2 stages (N = 1484), (F) N3 stages (N = 496). Abbreviations: relative (Rel.). (**C**) Sleep stages (first trace, top panel), instantaneous SB frequency (middle) and CA headpulse recording (bottom trace) from a 40-year-old female with no neurological disease. The CA recording is displayed with its DC components in the ±1g range in the vertical (Vert), anterior-posterior (AP), and lateral (Lat) planes. Across all subjects, SB frequency ebbed and flowed throughout the night, with a complete absence of SB at some points during the night. The highlighted timeframe (green region) in C is featured and expanded in D. (**D**) Runs of highly periodic SBs at a 0.04 bursts/min were observed in this subject (vertical dotted lines). Sleep bursts were mostly single bursts, like that shown in (A), but in this example of transition between N2 and N3 sleep, some were longer in duration. Fig. S2 shows an example where these “poly-bursts” were dominant.

Henceforth these forces are called “sleep bursts (SBs)”. These events are measurable using highly sensitive accelerometers attached to a headband worn during sleep (Fig. S1). SB appear to be generated by the brain (Figure 1B), and their dose is reproducible across devices buts changes with neurodegenerative disease (Table 2 and Figures 2 and 3). Due to the overlap in pulsatile frequency, we hypothesize that these SBs may reflect previously unobserved biological forces in humans associated with glymphatic drainage during sleep. This new biometric may serve as a valuable tool for future sleep research, especially in deepening our understanding of how sleep is restorative. It may also be useful for testing the hypothesis that neurodegenerative disorders are related to reduced glymphatic drainage because of the link between sleep loss and the development of these disorders.

**Figure 2.**
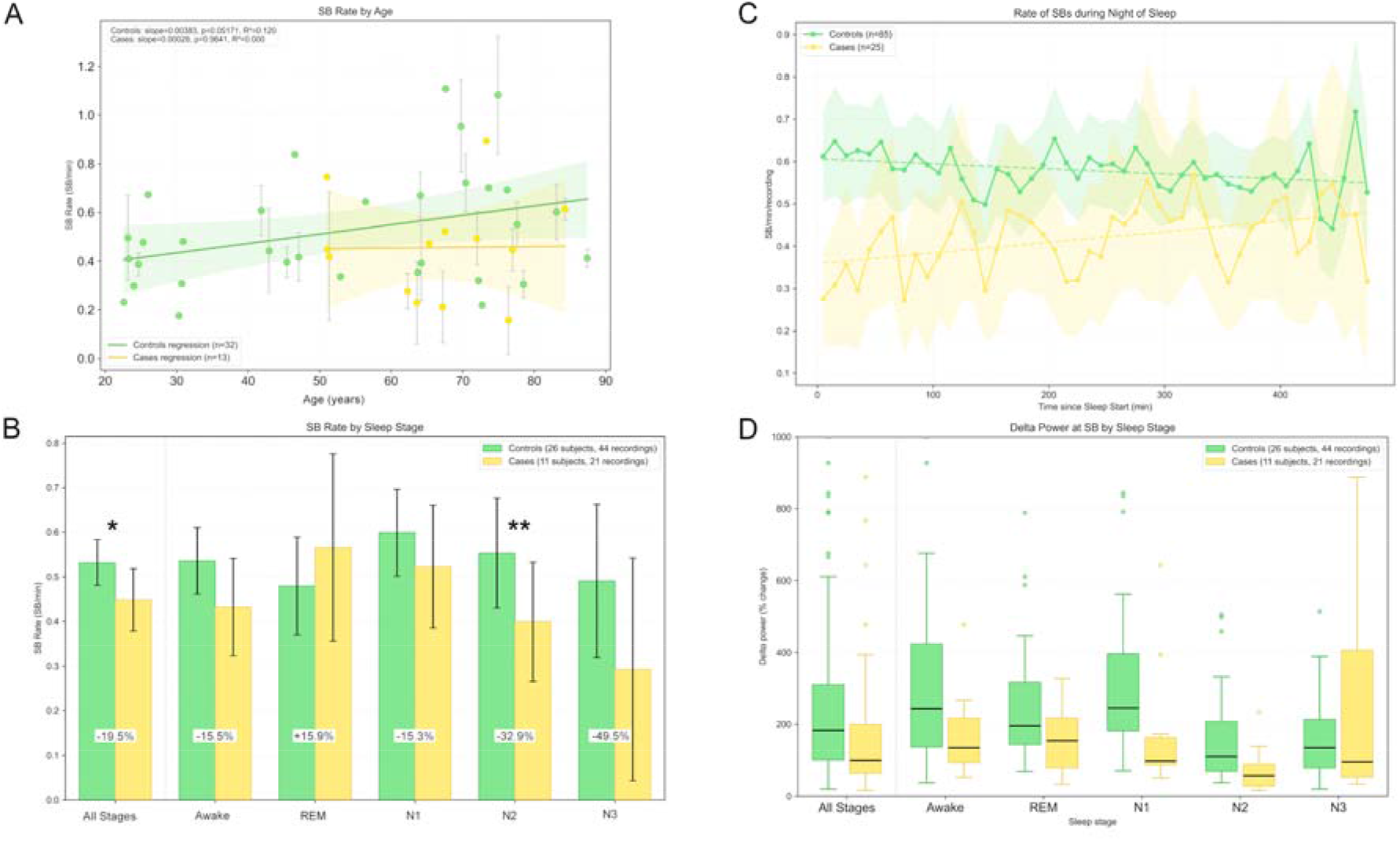
(**C**) SB rate was defined as the number of SB events over a specified time period. Control subjects and NDD subjects are plotted by their SB rate against age; subjects who had more than one night of sleep are shown with SD error bars and the mean SB rate was used in this analysis. There was a mild increase in SB rate with age in the control group (r^2^ = 0.12) and no correlation with age in NDD subjects. (**D**) SB rates compared by group (control or NDD cases) across sleep stages and all sleep. NDD cases had a significant 19.5% relative reduction in SB rates compared with controls (p = 0.041). There were also significant reductions in SB rate in N2 and N3 sleep in NDD subjects compared to controls. Comparing SB rates across each stage of sleep for controls found no significant differences by sleep stage including awake (p = 0.371, ANOVA). For NDD cases there was a trend toward lower SB rates toward deeper stages of NREM sleep (p = 0.091, ANOVA).

**Figure 3:**
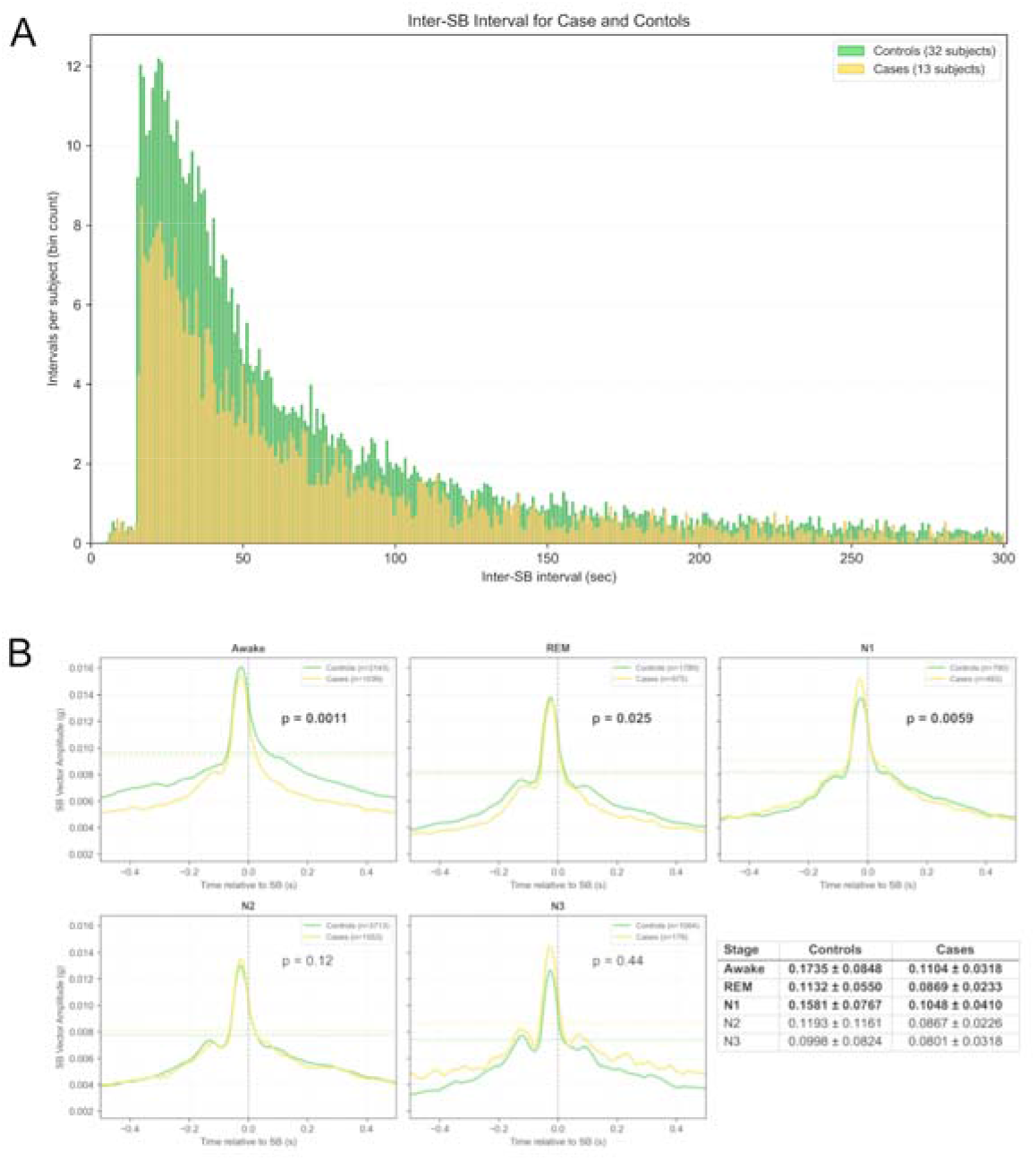
(A) Distribution of intervals between SBs in all subjects (total N = 24,837 SBs), with controls (green; N = 19,692 SBs) and cases (gold; N = 5,145 5 SBs). The distributions are not different in shape (p =0.43, Mann-Whitney test for location). The distribution is skewed with two preferred intervals: 3.75 bursts/min (0.0625 Hz) and 2.67 bursts/min (0.044 10 Hz). However, SB activity can be separated by multiple minutes (as shown out to 5 mins here). SBs occur at 0.54/min in controls and 0.42/min in cases because SB rate is summarized by totaling SB during the entire night of sleep and dividing by total sleep time. Examples of this more rapid recurrence of SB are shown in Figure 1 C,D and in supplemental Figure 2. The SB detection algorithm was made to be refractory for 15 seconds after finding the first rise of CA above threshold accounting for the fall off in intervals under 15 seconds. (B) SB triggered average of the rectified SB vector. Differences between controls and cases derived by measuring the full width at half maximum of the averages. Cases were significantly more narrow between Awake, REM and N1 sleep stages, and between N1-N3 stages combined (not shown) using Mann-Whitney U test. P values are shown for each comparison with significant differences in bold.

## Results

### Subjects

The occurrences of night-time SBs were assessed via accelerometry from a cranial accelerometer (CA) device worn during the night. Thirty-two volunteer subjects (“controls”, mean age 52.9 years, range [23-87 years], 13 female) agreed to wear the CA device during one or more nights of sleep (Table 1). Additionally, 13 subjects with NDD (“cases”, mean age 66.3 years, range [51-84], 4 female) had CA recordings. Concomitant ambulatory polysomnography using frontopolar EEG to identify wake and sleep stages was assessed in 26 controls and all 13 NDD subjects. There were no significant differences in median age, sex, or duration of sleep between controls with EEG recordings and those without.

**Table 1.**
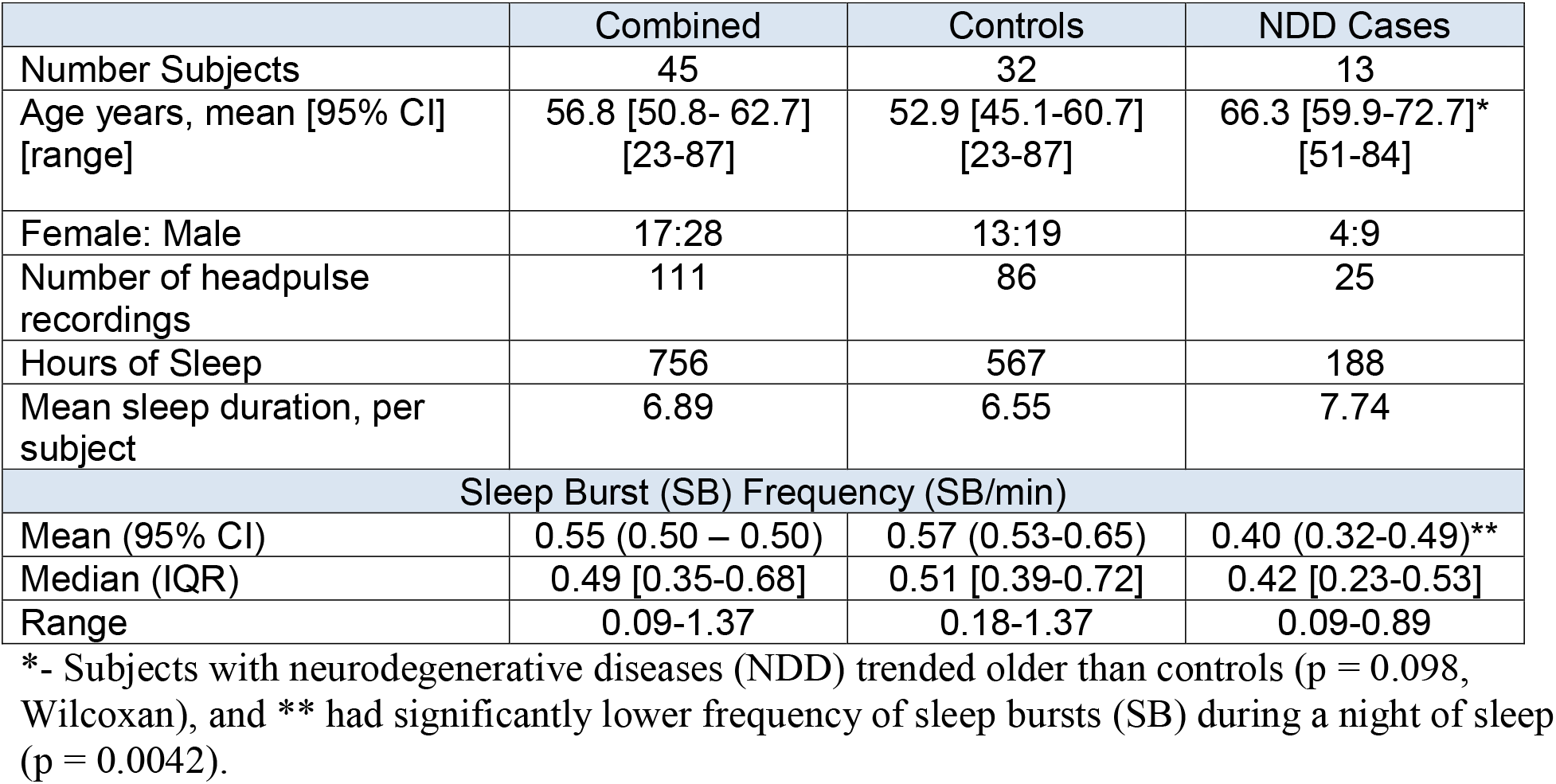
Demographics, Duration of Observation and Sleep Burst Frequency across recording arrangement.

|  | Combined | Controls | NDD Cases |
| --- | --- | --- | --- |
| Number Subjects | 45 | 32 | 13 |
| Age years, mean [95% CI]<br>[range] | 56.8 [50.8- 62.7]<br>[23-87] | 52.9 [45.1-60.7]<br>[23-87] | 66.3 [59.9-72.7]*<br>[51-84] |
| Female: Male | 17:28 | 13:19 | 4:9 |
| Number of headpulse recordings | 111 | 86 | 25 |
| Hours of Sleep | 756 | 567 | 188 |
| Mean sleep duration, per subject | 6.89 | 6.55 | 7.74 |
| Sleep Burst (SB) Frequency (SB/min) |  |  |  |
| Mean (95% CI) | 0.55 (0.50 – 0.50) | 0.57 (0.53-0.65) | 0.40 (0.32-0.49)** |
| Median (IQR) | 0.49 [0.35-0.68] | 0.51 [0.39-0.72] | 0.42 [0.23-0.53] |
| Range | 0.09-1.37 | 0.18-1.37 | 0.09-0.89 |

\*- Subjects with neurodegenerative diseases (NDD) trended older than controls ( $p = 0.098$ , Wilcoxon), and \*\* had significantly lower frequency of sleep bursts (SB) during a night of sleep ( $p = 0.0042$ ).

### Detecting Sleep Bursts

Figure 1A illustrates a typical sleeping CA measurement in a subject during stage N2 sleep. The normal “headpulse”(*11*) produced by cardiac contraction and detected by the CA device, appears as a time-varying signal of amplitude in the 10 – 30 milli-g range following each R-wave of the electrocardiogram (ECG) (Fig. 1Aa). Signal intensity varies slightly with respiration but typically follows a set pattern with each heartbeat(*11*). Superimposed on the normal headpulse, Figure 1Ab reveals three SBs as short-duration increases in cranial forces. Each SB was separated by 106 and 216 seconds in this example. The details of one of these SBs are shown in Figure 1Ac. SBs are distinguishable from the background headpulse by their larger amplitude (typically 2-5 times larger) with higher frequency components being temporally sharper compared to cardiac-produced headpulses. SBs are not correlated in timing with ECG R-waves (N=10 subjects with good quality ECG, CHI square goodness of fit P values ranged from 0.9998 to 1.0).

### Sleep Burst profiles

Examples of SB profiles are shown in Figures 1C and S2. A full night of headpulse recordings along with sleep staging is shown for a subject in Figure 1C. As was typical of all subjects, this individual had 1) periods of no SB present (Fig. 1C middle panel); and 2) a wide variance in SB frequency throughout the night. Some subjects had periods of periodicity (e.g., Fig. 1D) during the night. Another example shows a subject with highly periodic SBs (Fig. S2A), occurring approximately 33 seconds apart. A slow rise in CA magnitude appears to anticipate the SB (Fig. S2B), suggesting there is a biomechanical lead-up to the occurrence of a SB.

### Sleep bursts across age and sex

All 45 subjects had SBs across a total of 111 recordings comprising 756 hours of sleep (Table 1). Overall, SBs occurred at an average rate of 0.55 bursts/min. NDD subjects had a significantly lower SB rate of 0.40 SB/min compared to controls (0.57 SB/min, p = 0.0042). SB event frequencies, for each recording, are plotted by subject and age to illustrate the variance within and between individuals; SB frequency is not dependent on age (Fig. 2A). When accounting for multiple measures in subjects using a regression clustered analysis, SB frequency tended to increase with age in controls (p = 0.056, R^2^=0.117), but not in the NDD group. No sex differences in SB frequencies were found within groups (p = 0.47).

### Sleep bursts across sleep stages

The duration of sleep between cases and controls were not different except for an increase in REM duration in cases. Overall, SB frequency did not vary by sleep stage, or between wake after sleep onset and wake when lying, preparing to sleep (Table 2), though NDD trended towards a reduced rate during stages N2 and N3 (p = 0.091) (Table 2, Fig. 2B). In aggregate, 41 hours of wakefulness, 34 hours of REM, and 97 hours of NREM stages 1-3 (N1-N3) sleep were observed with simultaneous CA recordings. The durations and shapes of SBs were similar across subjects. The width at 50% maximum of rectified and averaged SBs were widest in Awake compared to REM, N2 and N1 sleep (P < 0.001), N1 wider than REM, N2 and N3 (p <0.04) (Table 2, Fig. S5). NDD subjects had significantly narrower SBs in Awake, REM and N1 sleep compared to controls (Fig S5). The temporal width of a SB is proportionate to biomechanical power.

**Table 2.** Sleep Burst Rates, Width and EEG Delta Power.

| Metric / Group | Awake | REM | N1 | N2 | N3 |
| --- | --- | --- | --- | --- | --- |
| **Mean SB Rate (SB/min)** |  |  |  |  |  |
| Controls | 0.53 | 0.49 | 0.61 | 0.54 | 0.46 |
| Cases | 0.45 | 0.57 | 0.51 | 0.36 | 0.23 |
| P (Welch) | **0.035** | 0.30 | 0.11 | **0.0025** | 0.063 |
| **SB width (sec)** |  |  |  |  |  |
| Controls | 0.174 | 0.113 | 0.158 | 0.119 | 0.0998 |
| Cases | 0.110 | 0.0869 | 0.105 | 0.0867 | 0.0801 |
| P (Welch) | **0.000054** | **0.011** | **0.00061** | 0.079 | 0.21 |
| **Relative Delta Power at SB (% change above baseline)** |  |  |  |  |  |
| Controls | 332 | 324 | 363 | 167 | 212 |
| Cases | 172 | 158 | 181 | 75.0 | 284 |
| P (Mann-Whitney) | **0.057** | 0.11 | **0.012** | **0.012** | 0.85 |

### Sleep burst frequency

SBs, when periodic, occurred at two modal frequencies: 3.75 burst/min and 2.67 bursts/min for all 20,449 SBs recorded (Fig. S4); these distributions were not different between cases and controls. Because there were periods during sleep where no SBs were observed (periods of cranial force quiescence), the frequency of SBs across the whole night of sleep is lower than these modal frequencies (average 0.55 bursts/min, Table 1). SB rates decline over a night of sleep for controls (Fig. 2C) but increase for NDD subjects. There is a significant difference in the SB rate in the first 30 minutes sleep between cases (0.63 SB/min) and controls (0.31 SB/min, p = 0.00041).

### Are sleep bursts associated with EEG changes?

To explore the possibility that SBs are generated by the brain, we tested the correlation between SB events and bifrontal EEG recordings. Signal averages of rectified EEG from 17 subjects, across all sleep stages, is shown in Figure 1B, with the center aligned to the SB event. Across all recordings, rectified EEG power rose ~ 1 sec prior to a SB event and persisted for 1-2 seconds following. Inspection of the stage-specific plots of EEG power by frequency band identified that in all sleep stages the delta frequency band is the largest contributor. The increase in power is predominantly in REM, N1 and N2 stages and least in N3 stages. The observation that EEG power changes prior to SB events suggests that brain activity is the driver of SBs. Delta power during a SB was lower in NDD subjects across all sleep stages (Fig 2D).

### Sleep bursts across headpulse device models

In participants who wore both headset models of the headpulse, CA devices (Fig. S1), the SB general phenomenology was stable within an individual across devices. More specifically, within an individual, SB width was stable, though the amplitude did vary. We accounted for these variations when detecting SB events by using the SB trace as described in the methods. The amplitude of the CA signal may be a function of the coupling of the device to the scalp so it may not be a reliable parameter.

### Are sleep bursts associated with periodic limb movements (PLMs)?

To test whether the short-duration forces detected on the head during sleep are produced from body movements, limb accelerometry was applied in 11 subjects, 3 of which reported a history PLMs. For the subject featured in Figure 1 with no reported PLMS, placement of accelerometers on all 4 limbs revealed transient forces that triggered the same detection algorithm for SBs. In this subject, 413 SBs were recorded, and 372 similar short-duration, low-amplitude forces in any limb were identified. Of all 785 events, 125 were within 0.5 seconds of each other. The probability this would occur by chance is low (p <0.00001, Ripley’s K function(*12*)). Importantly, the other 660 events were independent of each other, revealing that most SBs are unlikely to be due to limb movements. A similar result was observed in 7 other subjects without PLMS (including the subject shown in Fig. 2). Three subjects with reported PLMS underwent both head and leg measurements during a night of sleep. One subject with large amplitude PLMS demonstrated characteristic periodic leg movements at a frequency of 0.04-0.06 Hz throughout most of their sleep time (Fig. S3) which is similar in frequency to SBs. These forces were observed in the CA measurements and were of sufficient size to obscure smaller SBs and triggered the sleep burst detection algorithm. In this individual, SBs occurred 461 times and leg movements 633 times during the night. Of the 461 SBs captured, 154 of them were independent of any leg movements, revealing a greater overlap of leg movement events with SBs. Isolated SBs occurred within the same frequency range as the periodic leg movements. In two other subjects with PLMS, leg movements were independent from SBs. Inter-SB histograms of head and limb SBs are shown in supplemental figure S3.

## Discussion

We identified a previously undescribed, short-duration cranial-force phenomenon (“sleep bursts”) occurring during human sleep that can be measured noninvasively. The intermittent, increases of force on the human head occurred in all subjects assessed, across all sleep stages, exhibit periodicity and have characteristic short-duration morphology. The SB phenotype itself is altered in the presence of NDD. Specifically, in cases compared to controls, SB frequency is lower, SB width is narrower across all sleep stages. SB are temporally couple to changes in EEG activity, where the EEG power change precedes the mechanical event. This novel brain-associated biomechanical phenomenon of sleep has frequency and temporal characteristic changes which reduce the dose of SB experienced in the presence of neurodegenerative disease. This difference should be examined in future work as a biometric for the diagnosis of NDD.

### Sleep Bursts in the presence of periodic limb movements

SBs are not a phenomenon isolated to the head but occur in other body regions as well. Somewhere between 30-67% of the head and limb SBs were coincident, suggesting that overall, SBs are systemic and coordinated, but it is common for a body region to experience these forces independent of the entire body. The relative independence of SB in the limbs and head indicates that one is not passively causing the other. Based on the 3 individuals with known PLMS, SBs were not contaminated by body motion, indicating that SBs are not a result of PLMs. However, it does raise a question of whether PLMS share neurophysiology with SBs since they have highly similar frequency. The SB could result from a gross sympathetic discharge, resulting in changes in EEG power PLMs are known to be associated with concurrent EEG, cardiac, and autonomic events, and additional studies examining these concurrent events in persons manifesting both may help us to discriminate between them in a dispositive fashion(*13, 14*).

### Sleep Bursts are unlikely due to sleep apnea

Sleep apneic events are greater during REM sleep than other sleep stages and typically increase with age. SBs were measured in everyone assessed, independent of age or gender. Similarly, the frequency of SBs did not differ in REM sleep compared to other sleep stages. Therefore, it is unlikely that SBs are related to the occurrence of sleep apnea. While one individual assessed had reported sleep apnea, they used a CPAP during HP assessment and had similar SB profiles to the 31 subjects with no reported sleep apnea.

### Sleep Bursts may be a possible marker of glymphatic activity

SBs may be produced by the forces that drive glymphatic drainage in a pulsatile manner. Putatively, glymphatic clearance is associated with pulsatile activity due to cardiac contraction and pulsation within cerebral arterioles, respiration, and both low (0.023-0.73Hz) and very-low frequency (VLF, 0.001-0.023Hz) pressure waves in the perivascular spaces and interstitial fluid of the parenchyma, in the range that we detected here(*15*). In support of SBs possibly being due to glymphatic drainage, we note that whole-brain accelerations are produced by the arrival of blood ejected from the heart during cardiac contractions(*15*).

As proposed by Nedergaard and colleagues, the flow of glymphatic clearance may be through glial cells, facilitated by water channels in the periarteriolar space, and moved into the perivenular spaces of the brain(*7, 8*). This system of “check-valves” would facilitate flow toward the perivenular space if the pressure inside the parenchyma is increased. Flow through the brain interstitium has been hypothesized to be driven by the pulsatility of arterioles(*16*). Yet glymphatic drainage appears to be predominantly during sleep, and arteriolar pulsations are omnipresent. In our recordings, cardiac contractions were not associated with the SBs.

Vasomotor waves, occur in the VLF range and are driven by the sympathetic system. During sleep, VLF waves increase in power and peripheral VLF waves become more synchronized with central ones(*17*). Vasomotor activity, and vasoconstriction of the cerebral arterioles is regulated by the sympathetic system, which in turn is regulated by norepinephrinergic neurons in the locus coeruleus(*18*). In fact, periodic LC neuronal discharge generates the “vessel clench”-a change in vasomotion. LC stimulation has been associated with changes in vasomotion, which in turn was associated with the pumping action to drive CSF flow(*18, 19*); the change in CA size in a crescendo-decrescendo fashion (Figure S2) suggests a pumping action. Further, norepinephrine levels in the brain fluctuate at an approximate 50 s interval(*18*). Dual-photon imaging of cerebral arterioles in sleeping mice demonstrates periodic caliber changes occurring 1-2 times per minute(*19*).

Periodic, transient reductions in cerebral blood volume have been observed in human subjects sleeping in an MRI scanner as documented by periodic, transient reversal of CSF flow in the 4^th^ ventricle and cerebral aqueduct(*20*). This reversal lasts seconds and is likely caused by transient arteriolar vasoconstriction, making the intracranial intravascular blood volume transiently decrease. This transiently increases intraventricular CSF volume, as CSF briefly flows in reverse thought the 4^th^ ventricle. Once relaxation of cerebral vasoconstriction occurs, a transient increase in pressure gradient from brain to perivenular spaces arises, aiding the flow of interstitial and glial unidirectional flow of fluids into the perivenular space. Similarly, in Drosophila, hemolymph is moved through small parts of the body, particularly the antennae, not by heart contractions, but by the pulsatile contraction/relaxation of a muscle passing through the brain. The myogenic rhythm of the muscle was measured at 1.72 ± 0.4 Hz(*21*) and raised the proposition that brain pulsatility could aid in moving gas exchange in very small, remote parts of the brain. With the mechanism of transient increases in human CSF volume coupled with the return of normal cerebral blood volume, the brain should transiently increase pressure against the skull, which would be measurable on the skull surface. We propose that this as a possible source of SBs reported here.

### Sleep Burst Differences in Neurodegenerative Disease

Impaired glymphatic drainage has been hypothesized as enhancing the concentration of brain proteins aggregates found in various NDD (refs 7, 9, 10). Exploring this hypothesis requires a biometric of brain glymphatic activity. SBs may reflect ongoing glymphatic activity, and SB are lower in frequency and shorter in duration than in control subjects. This lower “dose” of SBs through a night of sleep would diminish brain cleansing and exacerbate neurodegeneration. Both the hypothesis of impaired glymphatic activity causing neurodegeneration and SBs as a measure of glymphatic drainage is the subject of ongoing research.

### Alternative sources of Sleep Bursts

SBs may alternatively represent microarousals, which are greater in N1, and can occur at a frequency of every 2-3 minutes, possibly regulated by LC discharge(*22*). However, SB frequency was independent of sleep stage. Deeper analysis of EEG features correlating with SB events is ongoing, however EEG delta power does increase before and following the SB supporting SB generation from a cerebrally mediated mechanism.

### Strengths and limitations of this research

There are several strengths of this research. Data were obtained from a broad range of ages (23-87 years old) in males and females free of sleep disorders (except three with PLMS and one with treated sleep apnea). Cases were well phenotyped through their involvement in UCSF Neuroscape and Fein Memory and Aging Center studies with a focus on neurodegenerative disease. Data were analyzed by a set algorithm, whereby SBs were identified by the same criteria in all subjects, making the documentation of an SB objective. No SB events were changed following algorithm identification. This likely led to some contamination of sleep-related movements in some SB identification, but the stereotyped shape and short duration of SBs across subjects argues against motion contamination. Frontal EEG was obtained in most subjects, providing data to correlate with sleep stage. We performed limb recordings in 11 subjects, showing that the SBs measured on the cranium are not due to gross body movements. There are weaknesses of this study. Although we did not observe a sleep-stage dependency for SBs, not all recordings were performed concomitant with frontal EEG. Also, we have not yet performed systematic analysis looking for SBs in awake subjects; it is difficult for subjects to remain still for long periods of time while awake. Further, two headpulse models were used that measured SBs, however, SBs were similar across both devices within a participant. Despite the highlighted weaknesses, we describe a novel, human, physiological measure needing further investigation.

## Acknowledgements

We thank all the subjects who participated in this study. Further, we thank the staff in both Dr. Wade Smith’s lab group and the Sleep in Aging and Neurodegenerative Disease lab group (Drs.Walsh & Neylan’s lab) at the University of California San Francisco. We particularly thank Macy Barros, Jonathan Shih, Quentin Coppola, and Felicia Song. We thank those who helped in referring/recruiting subjects and assisting with data collection, including Tanvi Thummala, members of UCSF Neuroscape, and members of the UCSF Fein Memory and Aging Center BrANCH, ALLFTD, and ADRC teams. We also thank those who read a draft of this manuscript, including Drs. Joaquin Anguera, Joseph Winer and Bryce Mander.

## Funding

The John Madden Family (WSS)

The Prospect Creek Foundation (WSS)

Rainwater Charitable Foundation (TCN, CMW)

National Institute of Health R01AG060477 (TN)

National Institute of Health R01AG064314 (TN)

## Author contributions

Conceptualization: CMW, TCN, PAL, WSS

Methodology: CMW, TCN, LY, NP, EDL, EL, JC, DR, SHW, PAL, WSS

Data Processing: LY, NP, EL, JC, WSS Analyses: WSS

Funding acquisition: CMW, TCN, WSS Writing – original draft: CMW, WSS

Writing – review & editing: CMW, TCN, LY, NP, EDL, EL, JC, DR, SHW, PAL, WSS

## Diversity, equity, ethics, and inclusion

Subjects provided informed consent prior to taking part in this study which went through ethical approval with the University of California San Francisco (UCSF) IRB. All cognitively healthy adults were eligible for this study. As a study team, we promote equity and diversity.

## Competing interests

CMW: Author CMW has been loaned Sleep Profiler devices from Advanced Brain Monitoring, Inc. for an unrelated study. CMW has co-authored papers with Advanced Brain Monitoring, Inc. CMW has received funding from the Rainwater Charitable Foundation as a member of the Tau Consortium, as well as funding from NIH. CMW co-founded and is an officer for a 501c3 nonprofit called the Advanced Science Exploratory Program. CMW declares that she has no other competing interests.

PAL: Author PAL holds stock as co-founder of MindRhythm, Inc.

LY: Author LY has provided consulting services to NextSense. This consulting work was unrelated to the current study and did not influence the design, analysis, or interpretation of the findings. LY declares that he has no other competing interests.

JC: The author JC declares that he has no competing interests.

NP: The author NP declares that she has no competing interests. EDL:

The author EDL declares that he has no competing interests. EL:

The author EL declares that she has no competing interests.

DR: Author DR declares that she has no competing interests.

SHW: Author SHW has been awarded patent USPTO 10,856,800. A related provisional patent application 37759_0690U1 has been filed under his name. Both inventions involve accelerometry of the head during sleep but use different methods to appose the sensor(s) to the head than the method described in this paper. Consistent with public law these are properties of the U.S. Department of Veterans Affairs where the inventor is employed and confer no financial benefits to him. Neither technology has been licensed.

TCN: Author TCN has been loaned Sleep Profiler devices from Advanced Brain Monitoring, Inc. for an unrelated study. TCN has co-authored papers with Advanced Brain Monitoring, Inc. TCN has received funding from the Rainwater Charitable Foundation as a member of the Tau Consortium, as well as funding from NIH. TCN declares that he has no other competing interests.

WSS: Author WSS holds stock as co-founder of MindRhythm, Inc.

## Data and materials availability

All data, code, and materials used in the analysis are available to any researcher for the purposes of reproducing or extending the analysis. A standard, materials transfer agreement will be used to share de-identified data in agreement with both UCSF’s data management policies and those of the requesting researcher’s institution.

## Supplementary Materials

Materials and Methods

Figs. S1 to S6

## Notes

### Summary of Updates

Revised authorship and modified the title from Sleep Bursts to Intermittent Cranial Accelerometric Pulses

## References

1. J. M. Siegel, Clues to the functions of mammalian sleep. Nature 437, 1264–1271 (2005).

2. K. C. Simon, L. Nadel, J. D. Payne, The functions of sleep: A cognitive neuroscience perspective. Proc Natl Acad Sci U S A 119, e2201795119 (2022).

3. M. P. Walker, R. Stickgold, Sleep-dependent learning and memory consolidation. Neuron 44, 121–133 (2004).

4. D. S. Ramanathan, T. Gulati, K. Ganguly, Sleep-Dependent Reactivation of Ensembles in Motor Cortex Promotes Skill Consolidation. PLoS Biol 13, e1002263 (2015).

5. F. Waters, V. Chiu, A. Atkinson, J. D. Blom, Severe Sleep Deprivation Causes Hallucinations and a Gradual Progression Toward Psychosis With Increasing Time Awake. Front Psychiatry 9, 303 (2018).

6. H. Bonilla-Jaime, H. Zeleke, A. Rojas, C. Espinosa-Garcia, Sleep Disruption Worsens Seizures: Neuroinflammation as a Potential Mechanistic Link. Int J Mol Sci 22, (2021).

7. J. J. Iliff et al., A paravascular pathway facilitates CSF flow through the brain parenchyma and the clearance of interstitial solutes, including amyloid beta. Sci Transl Med 4, 147ra111 (2012).

8. J. Herz, A. Louveau, S. Da Mesquita, J. Kipnis, Morphological and Functional Analysis of CNS-Associated Lymphatics. Methods Mol Biol 1846, 141–151 (2018).

9. N. A. Jessen, A. S. Munk, I. Lundgaard, M. Nedergaard, The Glymphatic System: A Beginner’s Guide. Neurochem Res 40, 2583–2599 (2015).

10. L. Xie et al., Sleep drives metabolite clearance from the adult brain. Science 342, 373–377 (2013).

11. W. S. K. Smith, KJ; Lovoi, P., Detection of a Novel Signal of Large Vessel Occlusion Stroke Using Cranial Accelerometry - The Headpulse. Stroke 50, (2019).

12. B. D. Ripley, The Second-Order Analysis of Stationary Point Processes. Journal of Applied Probability 13, 255–266 (1976).

13. F. Ferrillo et al., Changes in cerebral and autonomic activity heralding periodic limb movements in sleep. Sleep Med 5, 407–412 (2004).

14. M. Sieminski, J. Pyrzowski, M. Partinen, Periodic limb movements in sleep are followed by increases in EEG activity, blood pressure, and heart rate during sleep. Sleep Breath 21, 497–503 (2017).

15. V. Kiviniemi et al., Ultra-fast magnetic resonance encephalography of physiological brain activity - Glymphatic pulsation mechanisms? J Cereb Blood Flow Metab 36, 1033–1045 (2016).

16. J. J. Iliff et al., Cerebral arterial pulsation drives paravascular CSF-interstitial fluid exchange in the murine brain. J Neurosci 33, 18190–18199 (2013).

17. J. Tuunanen et al., Cardiovascular and vasomotor pulsations in the brain and periphery during awake and NREM sleep in a multimodal fMRI study. Front Neurosci 18, 1457732 (2024).

18. N. L. Hauglund et al., Norepinephrine-mediated slow vasomotion drives glymphatic clearance during sleep. Cell 188, 606–622 e617 (2025).

19. L. Bojarskaite et al., Sleep cycle-dependent vascular dynamics in male mice and the predicted effects on perivascular cerebrospinal fluid flow and solute transport. Nat Commun 14, 953 (2023).

20. N. E. Fultz et al., Coupled electrophysiological, hemodynamic, and cerebrospinal fluid oscillations in human sleep. Science 366, 628–631 (2019).

21. A. R. Kay, D. F. Eberl, J. W. Wang, Myogenic contraction of a somatic muscle powers rhythmic flow of hemolymph through Drosophila antennae and generates brain pulsations. J Exp Biol 224, (2021).

22. A. Luthi, M. Nedergaard, Anything but small: Microarousals stand at the crossroad between noradrenaline signaling and key sleep functions. Neuron 113, 509–523 (2025).

