## Supplemental Materials for "Intermittent Cranial Accelerometric Pulses - a Potential Marker of Neurodegenerative Disease"

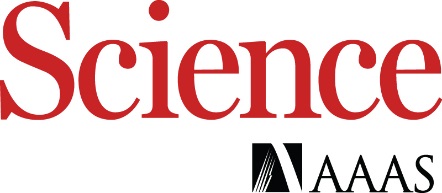

Supplementary Materials for

Intermittent Cranial Accelerometric Pulses - a Potential Marker of Neurodegenerative Disease

Christine M. Walsh^1^, Paul A. Lovoi^1^, Leslie Yack^2,3^, Joseph Chen^1,4^, Natalie Pandher^2^, Eliot D. Lee^1^, Thomas Hemphill^1^, Esther Li^2^, Dominica Randazzo^1^, Steven H. Woodward^5^, Thomas C. Neylan^1,2,3^, Wade S. Smith^1^*

**The PDF file includes:**

Materials and Methods

Supplementary Text

Figs. S1 to S5

References

**Other Supplementary Materials for this manuscript include the following:**

A. Materials and Methods

***Subjects***

Two cohorts of subjects were recruited. The first were “controls”, healthy individuals, who were free of a diagnosed neurodegenerative disease, stroke, and traumatic brain injury. The second group were “cases”, individuals identified as having mild cognitive impairment (MCI, n=8), Alzheimer’s Disease (AD, n=2), Progressive supranuclear palsy (PSP, n=2) or behavioral variant FTD (bvFTD, n=1). Subjects were consented to wear a commercial self-applied, frontopolar EEG device (Sleep Profiler, Advanced Brain Monitoring, Inc) and an experimental cranial-accelerometry headset designed to measure the headpulse (cranial forces) during one or multiple nights of regular sleep in their home environment. One subject had objective evidence of periodic limb movements (PLMs) from limb accelerometry during sleep. Another had a diagnosis of sleep apnea but was compliant with CPAP use. The study was approved by the University of California, San Francisco institutional review board (IRB).

***Methods***

*Sleep:* EEG was obtained using Sleep Profiler (Advanced Brain Monitoring, Carlsbad, CA). This device is an FDA-approved (Hardware: K152040; Software: K130007) frontopolar EEG (electrodes at AF7-AF8, AF7-Fpz, and AF8-Fpz positions) system with left and right electrooculograms (EOG), and submental electromyograms (EMG). Automated scoring was reviewed and edited by an investigator skilled in sleep stage scoring (LY) and was blinded to headpulse recordings. Sleep measures of interest were night-time wake, non-REM stage 1 (N1), non-REM stage 2 (N2), non-REM stage 3 (N3) and Rapid Eye Movement (REM) sleep.

*Headpulse:* The headpulse was measured continuously using two versions of a custom-built, battery-operated, non-invasive, Non-Significant Risk (NSR) device that transduced three-axis digital accelerometers with a least significant bit (LSB) sensitivity of 3.8 $\mu$g and a free scale range of $\pm$ 2 g. The first cranial accelerometer device included a plastic hairband with two printed circuit boards attached, each with an accelerometer (Fig. S1A). The dual, cranial accelerometer device was placed on the subject’s head in the coronal plane such that the accelerometers at the ends of the hairband lightly contacted the temporal bone just anterior to the tragus. This device also allowed transduction of the electrocardiogram. The second device was a smaller, single cranial accelerometer version (Fig. S1B) that attached via hook and loop fastener to a commercially available, adjustable elastic headband (Frog Sac, Amazon) or via hook and loop fastener to the Sleep Profiler, which used a hook and loop fastener that allowed adjustment to fit comfortably while sleeping and was positioned over the forehead.

Cranial accelerometry signals were digitized at a resolution of 20 bits and sampled at a rate of 250 samples per second. Data were stored on a micro-SD card within the headset for off-line analysis. Data were imported from the memory card and displayed using custom software written in Python V3.12. Both devices were synchronized by their internal real-time clocks, and minor adjustments were made to align the waveforms by changes in body position using custom software.

Head movements caused by change in head position, or awake epochs where the patient ambulated, produced g-forces exceeding 30 milli-g (gross body motion, GBM), which could be easily identified by amplitude and were excluded from analysis using an algorithm. Headpulse amplitude free of gross body motion is typically in the 2 – 15 milli-g range. Periodic, to semi-periodic increases in amplitude were discovered by visual inspection of accelerometry traces during human sleep; the first observed case is shown in Fig. 2B. We termed these transient or short-duration increases in forces that stood out over the headpulse background as a “Sleep Burst” (SB) (Fig. S2). We created an automated process in software to identify SB within and across subjects to provide a consistent method for studying SBs. The automated process of detection employed construction of a “sleep burst trace”. This was constructed first by normalizing each accelerometer signal to the root-mean square value of a moving sample $\pm$ 3 s from the central bin to eliminate changes in amplitude by varying forces of the device to the scalp as would occur when changing body position. Then the maximum absolute value of each of the three signals was constructed bin-for-bin to produce the SB trace. Shown for example in Fig. S2C, the SB trace amplifies transient, brief increases in the headpulse signal over the background headpulse. A SB was defined as a signal that rose typically above 7 in the sleep burst trace (unitless scale); this threshold was chosen to maximize detection by minimizing falsely identified gross motion masquerading as sleep bursts. Once the onset was identified, the algorithm waited at least 15 seconds before detecting a new sleep-burst trace. By visual inspection most SBs lasted under 2-3 seconds, but some could have multiple rising edges in one “poly burst”. This was inconstant across subjects, so we chose a refractory period of 15 seconds to homogenize the reporting of SB rates for the purpose of this study.

***Combining headpulse and sleep metrics*:** Using EEG and headpulse merged data, the number of SBs was determined and assigned one of five defined stages: Awake, REM, N1, N2, and N3 sleep stages. The number of SBs were reported as a frequency during each sleep stage in SB/min. The width of the SB signal was calculated by signal-averaging the sleep burst trace during each stage of sleep. The width was defined as the width of the averaged signal at the 50^th^ percentile of height (“Full Width at Half Maximum, FWHM). The periodic frequency of SBs independent of sleep stages was reported by the mean reciprocal of inter-sleep burst events. In cases where the headpulse was measured without EEG, SBs were identified and counted but were not analyzed by sleep stage. Sleep Start (SS) and Sleep Finish (SF) were identified by polysomnography in subjects who wore the Sleep Profiler. Bedtime (BT) was manually assigned at a time following the start of a recording when clear body motion was detected in the accelerometers preceding a period of inactivity. For the CA recordings done without the Sleep Profiler, BT, SS and SF were manually assigned in the order of SS – where the patient became motionless for several minutes, SF when the patient began to move in a sustained fashion that was not followed by inactivity, and BT was determined in the same fashion as with the Sleep Profiler subjects.

***EEG Analysis***: Time-frequency analysis of the single EEG electrode was performed using *FieldTrip*(*24*)*.* Preprocessing steps included band-pass filtering between 0.3 – 35 Hz and manual ocular artefact rejection. Time-frequency decomposition was performed using sliding-window Fourier transforms with a single Hanning taper. Frequencies from 0.5 to 30 Hz were analyzed in 0.05 Hz increments using adaptive window lengths of 3 cycles per frequency of interest across -5 to 5 seconds relative to sleep burst onset in 10 ms steps. Data were zero-padded to the next power of 2. Baseline values at -5s to -4s relative to sleep burst onset was used to calculate relative change and plotted with cool heatmap colors denoting reduced spectral power and warm heatmap colors denoting increased spectral power. Rectified, averaged, EEG signals from -5 to 5 s were also plotted. To calculate relative delta power for each SB, EEG was extracted in a ±10 s window centered on SB onset and analyzed with a short-time Fourier transform. Instantaneous delta-band power (0.5–4 Hz) was obtained by integrating the squared spectral amplitude over that frequency range at each time point. Relative delta power was then computed as the percent change from a within-sweep baseline defined as the average of the left and right ends of the window. The SB-associated value for each recording and sleep stage was the maximum of this relative series within the window; multi-night values were averaged within subject and stage before group summaries.

***Head and limb recordings:*** To assess for the contribution of periodic limb movements (PLMs) to SBs, participants were asked to place the accelerometer on both the head and ankle for a night of recording during sleep. The precise timing of gross leg movements and SBs were then assessed. Concomitance was calculated by finding all SBs that had a leg-associated SB within 3 seconds (6-second window). Six seconds was used based on the average duration of a leg sleep burst. An SB was considered “isolated” if an SB in the alternate measurement did not fall within this 6-second window.

For $O_{c}$ = observed number of coincident sleep bursts, $O_{nc}$= number of non-coincident sleep bursts, $n_{h}$= number of sleep bursts recorded from the head, $n_{p}$= number of sleep bursts recorded from the periphery, the overall Chi-square statistic ($x_{Overall}^{2}$) to reject the null hypothesis (that the SBs are occurring independent of each other) is:

$$x_{c}^{2}=\frac{\left( O_{c}- E_{c} \right)^{2}}{E_{c}}$$

$$x_{nc}^{2}=\frac{\left( O_{nc}- E_{nc} \right)^{2}}{E_{nc}}$$

$$x_{Overall}^{2}= x_{c}^{2}+ x_{nc}^{2}$$

The expected number of coincident SBs assuming the null hypothesis ($E_{c})$ can be calculated for bursts happening in a 6-second window, over the duration $D$ (seconds) of a recording as follows:

$$P\left( h\cap p \right)=P\left( h \right)P\left( p \right)$$

$$=\left( \frac{n_{h}}{D} \right)\left( \frac{n_{p}}{D} \right)$$

$$E_{c}=P\left( h \cap p \right)\frac{D}{6}= \frac{n_{h}n_{p}}{6D}$$

The expected number of non-coincident bursts assuming the null hypothesis ($E_{nc}$) is:

$E_{nc}= n_{h}+ n_{p}- E_{c}$

***Statistics:*** Correlation between electrocardiogram R-waves was tested by performing a histogram of SB to R-wave latencies between +/- 1 minute, then calculating the Chi-square goodness of fit for uniformity and p-values reported based on the degrees of freedom for all subjects with quality ECG tracings. As an additional test for the relationship between the sleep bursts measured on the head and the limb cluster, the Ripley K formula was applied.

B. Headset Photos

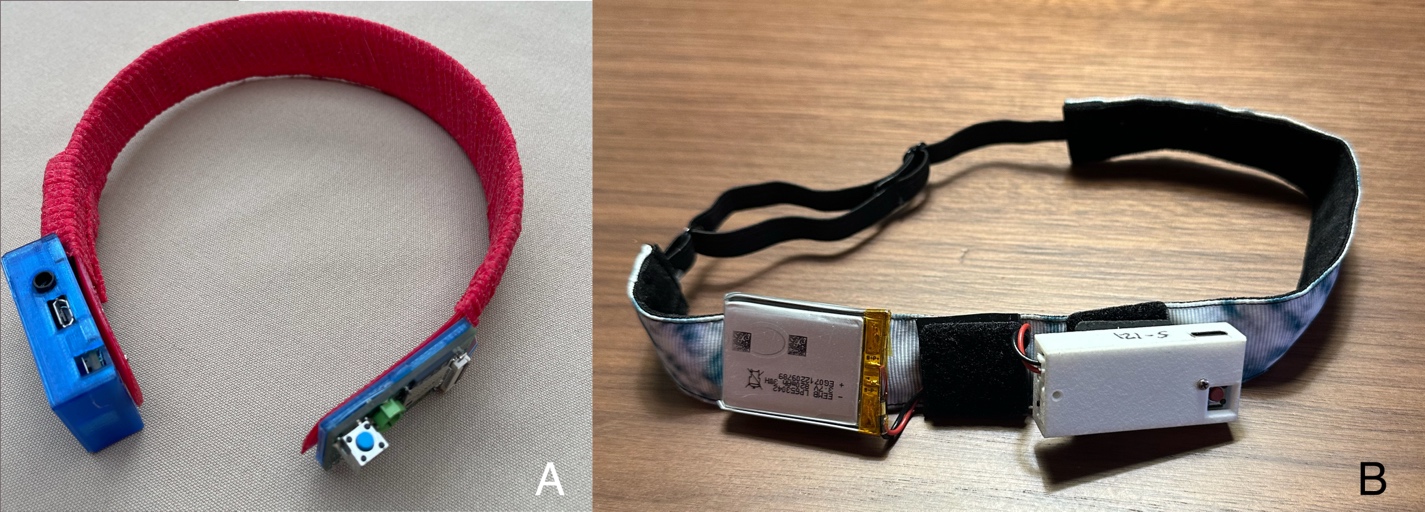

**Figure S1**. Custom headsets used in this study. (A) Two circuit boards were secured to a simple plastic headband and connected by a ribbon cable. The right side of the headset (with plastic cap removed) houses a microprocessor and micro-SD card holder for data storage along with a 3-axis accelerometer. The left side has another accelerometer along with battery charging circuitry and clock. The device connects via Bluetooth to a custom iPhone App written to start and stop recordings. (B) Smaller single channel version using a soft head strap with the device attached via hook and loop fastener. This device is placed on the forehead and is more comfortable to wear during sleep.

C. Sleep Burst Periodicity and Presence of Poly-Bursts

The periodicity of SB occurrence ranged from highly periodic (as shown in Figure S3 below), to semi-periodic (Figure 1 main paper). The subject below discovered this signal. Narrow sleep bursts begin just before sleep onset, then occur every 30-50 seconds. As sleep progresses, the bursts are accompanied by several secondary narrow SBs forming a “poly burst”. No other subjects had a mostly poly-bursts. The magnitude of the headpulse signal between SB events is suppressed following a SB event, then rises until the next SB potentially reflecting a change in intracranial compliance during the cycle. Other normal older adults did not exhibit multiple poly-bursts.

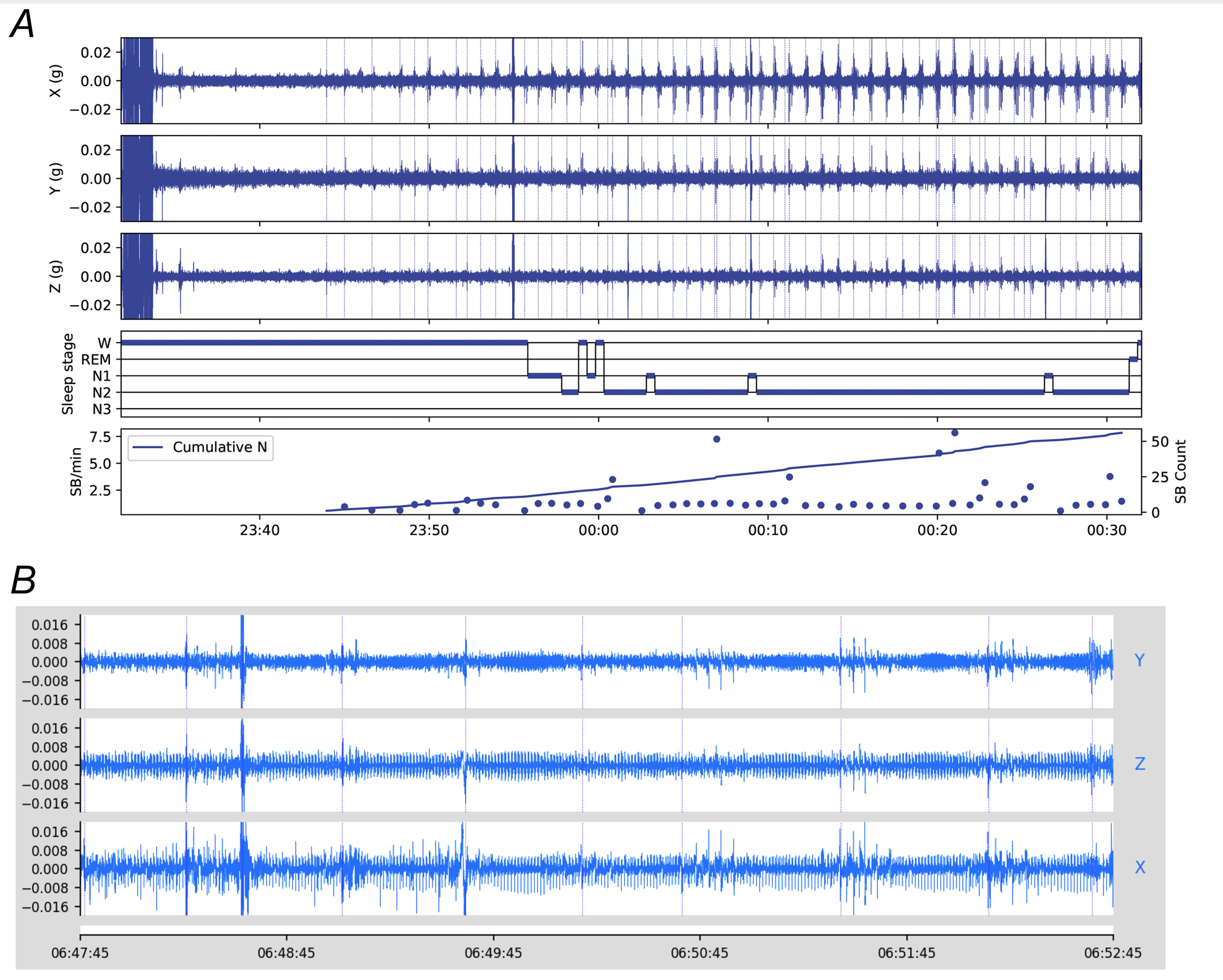

**Figure S2:** (A) First hour of sleep in a 72-year-old male subject. Sleep bursts began prior to polysomnographic detection of sleep onset, then began to build in amplitude with the addition of multiple brief spikes (poly-spikes) in the CA amplitude. The frequency of these SBs was highly regular, happening at 2 SB/min (0.033 Hz). X-axis is clock time. SB are denoted by vertical dotted lines. (B) An expanded view of a different night of sleep in the same subject shows the morphology of these poly-spike events over a 5-minute period.

D. Periodic Limb Movements and SB coincidence

These are supplemental figures for the main manuscript concerning the coincidence of limb SB and head SB.

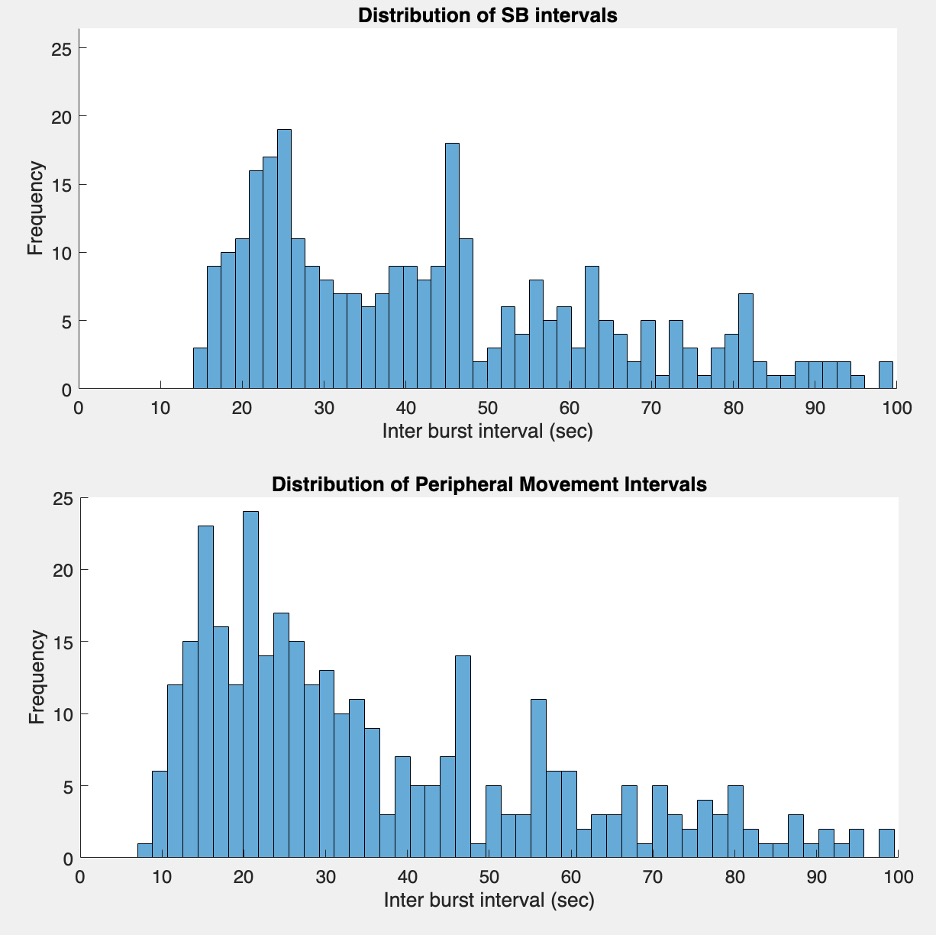

**Figure S3:** Top histogram- distribution of SBs in a subject with a diagnosis of PLMS; modal frequency is 2.4/min. Bottom histogram- distribution of left leg movements during the same night of sleep; modal frequency is 2.9/min. A chi-square test comparing the binned distributions of inter-burst intervals for SB events and peripheral movements revealed no significant difference between them (χ²(91) = 93.8, p = 0.4004).

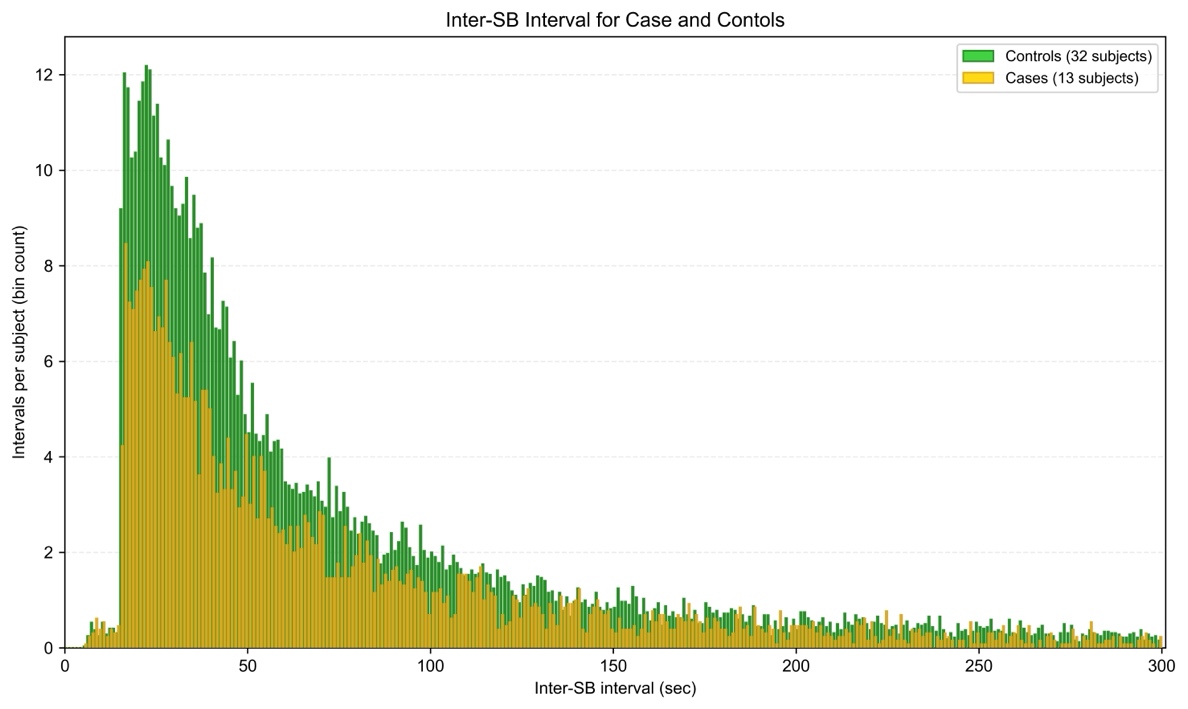

**Figure S4:** Distribution of intervals between SBs in all subjects (total N = 24,837 SBs), with controls (green; N = 19,692 SBs) and cases (gold; N = 5,145 SBs). The distributions are not different in shape (p =0.43, Mann-Whitney test for location). The distribution is skewed with two preferred intervals: 3.75 bursts/min (0.0625 Hz) and 2.67 bursts/min (0.044 Hz). However, SB activity can be separated by multiple minutes (as shown out to 5 mins here). SBs occur at 0.54/min in controls and 0.42/min in cases because SB rate is summarized by totaling SB during the entire night of sleep and dividing by total sleep time. Examples of this more rapid recurrence of SB are shown in Figure 1 C,D and in supplemental figure 2. The SB detection algorithm was made to be refractory for 15 seconds after finding the first rise of CA above threshold accounting for the fall off in intervals under 15 seconds.

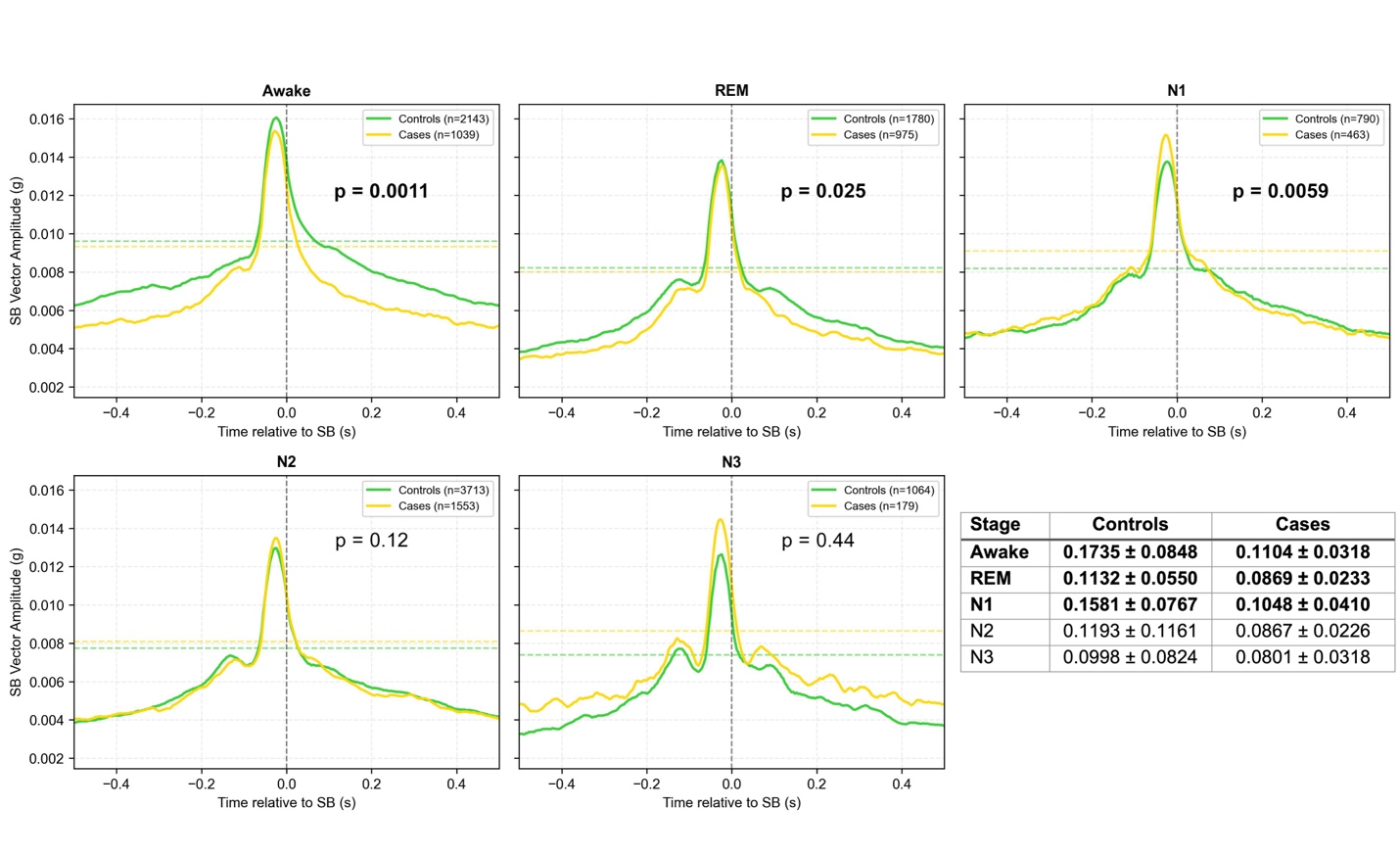
**Figure S5:** SB triggered average of the rectified SB vector. Differences between controls and cases derived by measuring the full width at half maximum of the averages. Cases were significantly more narrow between Awake, REM and N1 sleep stages, and between N1-N3 stages combined (not shown) using Mann-Whitney U test. P values are shown for each comparison with significant differences in bold.

*

*

**Figure S6:** Mean duration of controls and cases by sleep stage. * Compared to controls, N1 and N2 average durations were significantly longer for NDD subjects (N1 p = 0.046, N2 p = 0.015, Mann-Whitney U test).
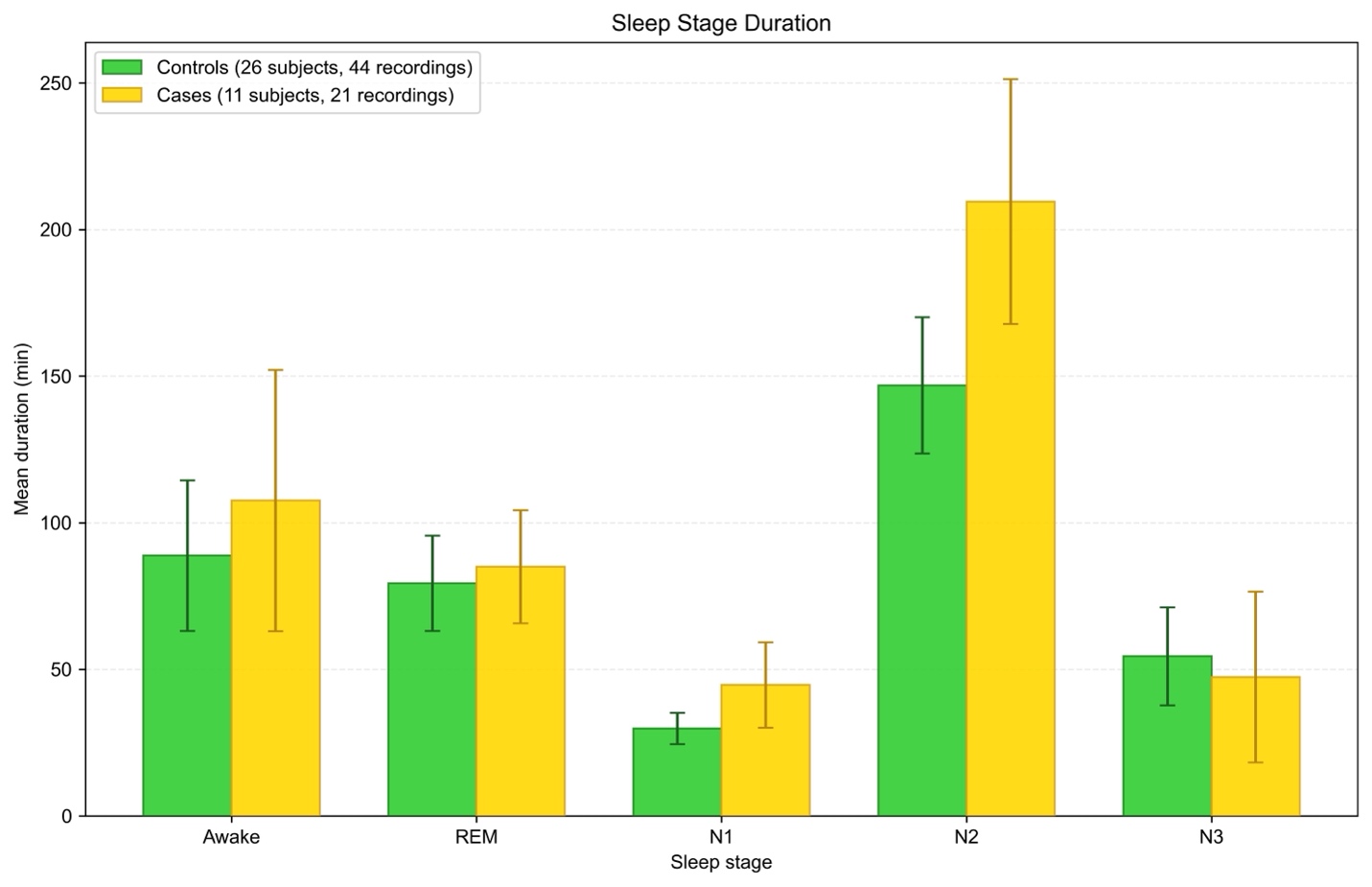
